# Evidence for domain-general arousal from semantic and neuroimaging meta-analyses reconciles opposing views on arousal

**DOI:** 10.1101/2024.05.27.594944

**Authors:** Magdalena Sabat, Charles de Dampierre, Catherine Tallon-Baudry

## Abstract

The term arousal is very often used, but classical textbooks from different domains of neuroscience and psychology offer surprisingly different views on what arousal is. The huge number of scientific articles with the term arousal (∼50.000) highlights the importance of the concept but also explains why such a vast literature has never been systematically reviewed so far. Here, we leverage the tools of natural language processing to probe the nature of arousal in a data-driven, comprehensive manner. We show that arousal comes in seven varieties: cognitive, emotional, physiological, sexual, related to stress disorders, to sleep, or to sleep disorders. We then ask whether domain-general arousal exists, and run meta-analyses of the brain imaging literature to reveal that all varieties of arousal, except arousal in sleep disorders for lack of data, converge onto a cortical arousal network composed of the pre-supplementary motor area and the left and right dorsal anterior insula. More precisely, we find that activity in dysgranular insular area 7, the region with the highest convergence across varieties of arousal is also specific to arousal. Our results show that arousal corresponds to a construct at least partially shared across different domains of neuroscience and identify the domain-general cortical arousal network. Novel taxonomies of arousal reconciling seemingly opposing views on what arousal is should thus include domain-general arousal as a central component.

**Significance statement:** The term arousal has been used in almost 50.000 scientific papers, but it is only loosely defined. The few attempts at defining arousal in neuroscience and psychology resulted in divergent views: arousal as a multi-dimensional construct or arousal as a global state. Is arousal an abstraction of the scientists’ mind reflecting a myriad of distinct processes, or is there some common neuronal feature? We used large-scale semi-automatic text mining methods and neuroimaging meta-analyses to review this vast and heterogeneous literature. Our results reveal the existence of domain-general arousal, a process shared by situations as different as a cognitive task, an emotional context, and the transition to wakefulness or sexual behavior. Domain-general arousal reconciles the concepts of general and multi-dimensional arousal.

## Introduction

Arousal is very often put forward as an explanatory variable for neuronal activity, bodily physiological parameters, and behavior in basic, translational, and clinical research, in both humans and animals. Given the importance of the concept, there has been surprisingly few attempts at establishing a taxonomy of arousal. Textbooks^1,2^ agree that arousal refers to brain-body states related to wakefulness and fluctuations within the wake state, that arousal modifies both central and autonomic nervous system activity, and that it is expressed in a myriad of bodily parameters. However, textbooks disagree on whether arousal is a unitary concept. On the one hand, in *Principles of Neural Science* (2021, p.1025), arousal is defined as a general, global state: “ […] a general state of arousal, which ranges from excitement and vigilance to drowsiness and stupor”. Arousal can indeed appear as a global factor organizing behavior. The Yerkes-Dodson law^3,4^ states that both low and high arousal drive performance down, while intermediate arousal corresponds to optimal performance. The prevalence of the inverted-U-shape relationship between arousal and behavioral performance across many different sensory and cognitive tasks in both humans and animals suggests some underlying common neural principle^5^. On the other hand, as advocated in the *Handbook of Psychophysiology* (2017, p. 415), the notion of general arousal should be rejected: “data […] were indeed incongruent with the idea that arousal is a unitary process”, and “arousal likewise became understood as a multidimensional process or construct”. Indeed, arousal arises from different ascending neuromodulatory systems (e.g., cholinergic, noradrenergic, etc.), it translates into different physiological changes (e.g., pupil diameter, heart-rate, skin conductance, etc.) that sometimes correlate only weakly with each other, or with subjective arousal ratings^5^.

Reconciling such divergent views calls for refining definitions and concepts to establish a taxonomy of arousal. Here, we probe for the existence of domain-general arousal. Do different varieties of arousal share a common neuronal basis? And even before this, how many varieties of arousal can be identified in the literature? Answering these questions requires analyzing the almost 50.000 scientific articles mentioning arousal that have been published as of April 2023. A few attempts at synthetizing the literature pioneered the field^6,7^, but their scope was necessarily limited by *a priori* selections of both articles and varieties of arousal. Here, inspired by recent work on data-driven cognitive ontologies^8–10^, we leverage the tools of natural language processing^11^ and graph theory to comprehensively analyze this large and heterogeneous literature. We identify in the literature seven different varieties of arousal in a purely data-driven manner. We then conduct meta-analyses^12,13^ of the brain imaging literature in humans and find that different varieties of arousal share common cortical resources.

## Results

### Identifying domains of the arousal-related literature using semantic analysis

#### The arousal literature is organized in seven semantic communities corresponding to known fields of research

We retrieved the 49 525 scientific references returned by the exact query “arousal” from Web of Science and PubMed and show that those articles are organized in seven meaningful semantic communities (Figure 1). More precisely, from the 49 525 abstracts we automatically extracted terms, i.e., associations of two to three words, and selected the most frequent ones. We then excluded non-specific terms (e.g., “significant result” or “previous research”, see Supplementary Table 1), as well as all brain-related terms (e.g., “brain imaging” or “prefrontal cortex”), and finally retained the remaining 1200 most frequent terms to build a graph reflecting the semantic content of the arousal literature. In this graph, terms are node, connected by edges that reflect how often terms co-occur in the references.

**Figure 1.**
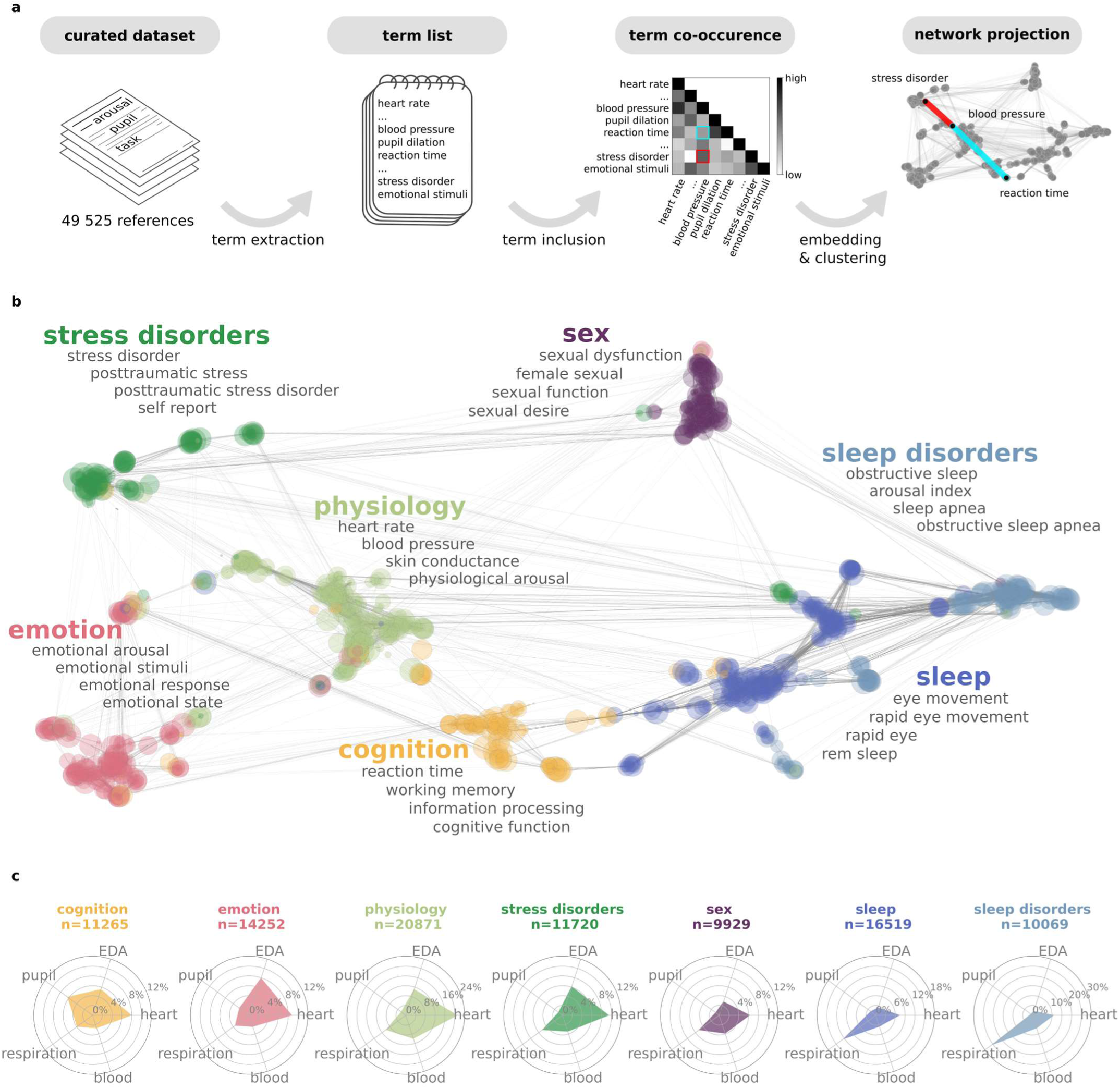
The semantic graph of arousal-related literature. A. Schematic description of the semantic analysis workflow. From a dataset of 49 525 references containing the word “arousal”, we extracted the terms (association of two or three words) that appeared most frequently and computed the co-occurrence of these terms. Subsequently, the co-occurrence matrix is embedded into a semantic graph. In the semantic graph, terms that tend to co-occur, and to co-occur with the same other terms, are closer, forming a semantic community. B. The semantic graph of the arousal literature constructed on the 1200 most frequent terms. Each dot corresponds to a term, each edge corresponds to semantic similarity. Nodes that cluster together are assigned the same color. The four most frequent terms are listed for each community in gray font and were used to assign a label to each community (color, bold font). C. Physiological profiles of the semantic communities, showing the percentage of articles within each semantic community using terms related to eye-tracking/pupillary (label: pupil), cardiac (heart), respiratory (respiration), blood pressure (blood) or skin conductance (EDA) measures of arousal. *Abbreviations:* EDA-electrodermal activity.

The semantic graph of the arousal literature reveals seven communities (Figure 1B) that we associate with seven varieties of arousal: arousal in cognition, emotion, physiology, stress disorders, sex, sleep, and sleep disorders. This association is based on the most frequent terms (as illustrated in Figure 1B) as well as on the terms with the highest number of connections within each semantic community (Supplementary Table 2).While we illustrate the results obtained with the analysis of the 1200 most frequent terms, the seven communities and their respective locations in the graph are stable across a wide range of number of terms included in the analysis (from 300 to 2285, Supplementary Figure 1). The physiological arousal community is located at the center of the overall semantic space, the two arousal in sleep communities are close together, arousal in stress disorders sits next to emotional arousal, and sexual arousal is far away from all other semantic varieties of arousal.

#### Each semantic community has a specific physiological measure profile

To cross-validate the existence of seven different varieties of arousal as identified by the semantic graph, we reasoned that if the semantic communities indeed refer to distinct fields of research, then they should differ in the measures used to operationalize arousal. We thus quantified the representation of physiological measures in the articles contained in each semantic community. We first assigned each article to semantic communities: if a term from a given community was indexed in the abstract, the article was considered belonging to this community. An article might thus project onto more than one semantic community. We inspected the occurrence of the words related to five common physiological measurements used as arousal proxies: cardiac activity, eye-tracking, respiration, skin conductance and blood pressure (dictionary of terms in Supplementary Table 3).

The resulting “physiological profiles” (Figure 1C) constitute distinctive signatures of each semantic community. Measures of respiration dominate in the two sleep communities, cardiac measures are dominant in the stress and physiology communities, while pupillometry is almost exclusively used for the study of cognitive arousal. Altogether, each arousal community uses a specific combination of bodily measures of arousal. Interestingly, although pupillometry is an increasingly popular measure of arousal especially in animal studies^14^, eye-tracking is not a dominant one in any of the communities. It is worth underlining that we obtain this result even though an article and its associated physiological measures could be assigned to more than one semantic community. In other words, across fields of research, arousal is associated with distinct physiological profiles. This result cross-validates our approach of categorizing arousal literature, as it retrieves field-specific experimental preferences.

### Neuroimaging meta-analysis: identifying brain regions associated with arousal across fields of research

#### The arousal cortical network: three cortical regions where different varieties of arousal converge

The semantic analysis reveals that arousal is a term employed in fields that differ by their object of study (i.e., sleep vs. emotion) and in how they measure arousal (i.e., pupil diameter vs. respiration). Is there nevertheless a shared neuronal substrate for arousal, common to the seven varieties of arousal we identified, or are the varieties of arousal so different that they share no common neuronal substrate? To answer this question, we first ran, for each field of research separately, an activation-likelihood estimation analysis on 415 neuroimaging studies available in the open-source NeuroQuery^15^ database. We excluded studies using a priori regions of interest, since regions of interest constitute a known biasing factor in meta-analysis^16,17^. We based our Activation Likelihood Estimation (ALE)^12,13^ meta-analysis on the remaining 228 studies, of which 221 ( corresponding to an estimate of 6647 participants) could be reliably assigned to at least one semantic community when using 1200 terms to generate the semantic graph. Except for sleep disorders, where the number of available studies (n=6) is very low, we can identify a network associated with arousal in each field of research, as shown in Figure 2. Some arousal-related regions are found selectively in some fields of research but not others – for instance the left lateral occipital cortex is consistently found in emotional and sexual arousal but is not observed for other varieties of arousal. Still, there is convergence of four or five varieties of arousal in several regions.

**Figure 2.**
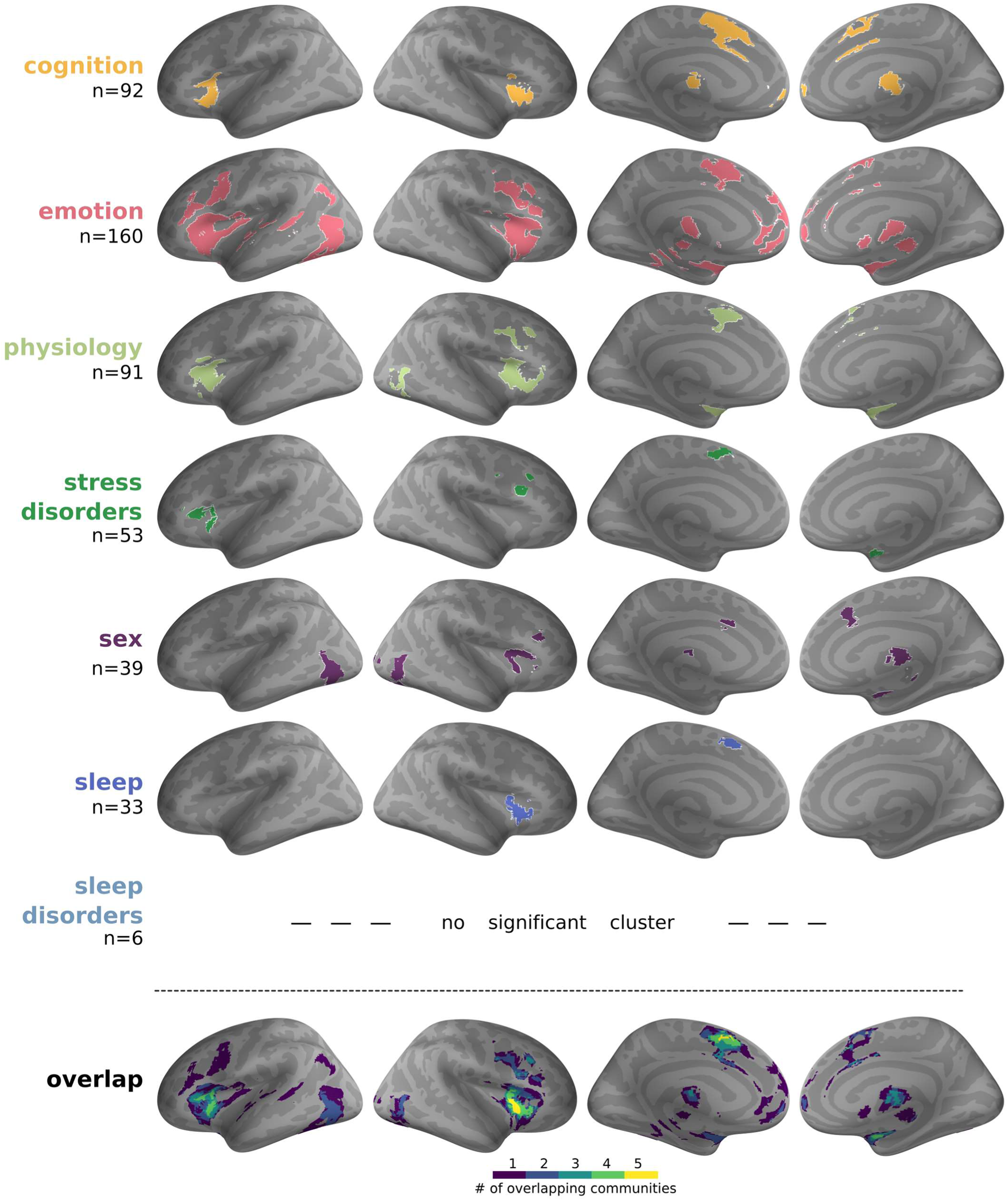
Meta-analysis of arousal in the seven varieties or arousal and their overlap. Results of the ALE meta-analysis for each semantic community (semantic analysis with 1200 terms), projected for visualization on inflated surfaces (voxel-level cluster forming *p* <0.001 uncorrected; cluster level *p* <0.05 corrected for multiple comparisons using Monte-Carlo permutations). The ‘n’ indicates the number of fMRI studies included in each meta-analysis. Note that because an article could be assigned to more than one semantic community, the sum of the number of fMRI studies included in each meta-analysis is larger than 221. The number of available imaging studies for the sleep disorders community (n=6) was too low to obtain significant results. The bottom row shows the overlap across activations obtained for each variety of arousal.

Our results reveal the existence of a cortical arousal network, composed of three regions where four or five different varieties of arousal converge (Figure 2, bottom row): the right and left anterior insula, and pre-supplementary motor areas (preSMA). As detailed in Table 1, cognitive, emotional and physiological arousal converge on all three nodes of the cortical arousal network, but left and right anterior insula differ when it comes to other types of arousal: sexual arousal and arousal in sleep appear in the right anterior insula, and arousal in stress disorders in the left anterior insula. Sexual arousal is absent from preSMA, and arousal in stress disorders and arousal in sleep overlap only marginally within this node, arousal in sleep being more anterior than arousal in stress disorders in preSMA. These results hold true even when considering unthresholded meta-analytical results (Supplementary Figure 2). In other words, all six varieties of arousal for which enough data exist load, sometimes with different weights, on the three nodes of the cortical arousal network.

**Table 1.**
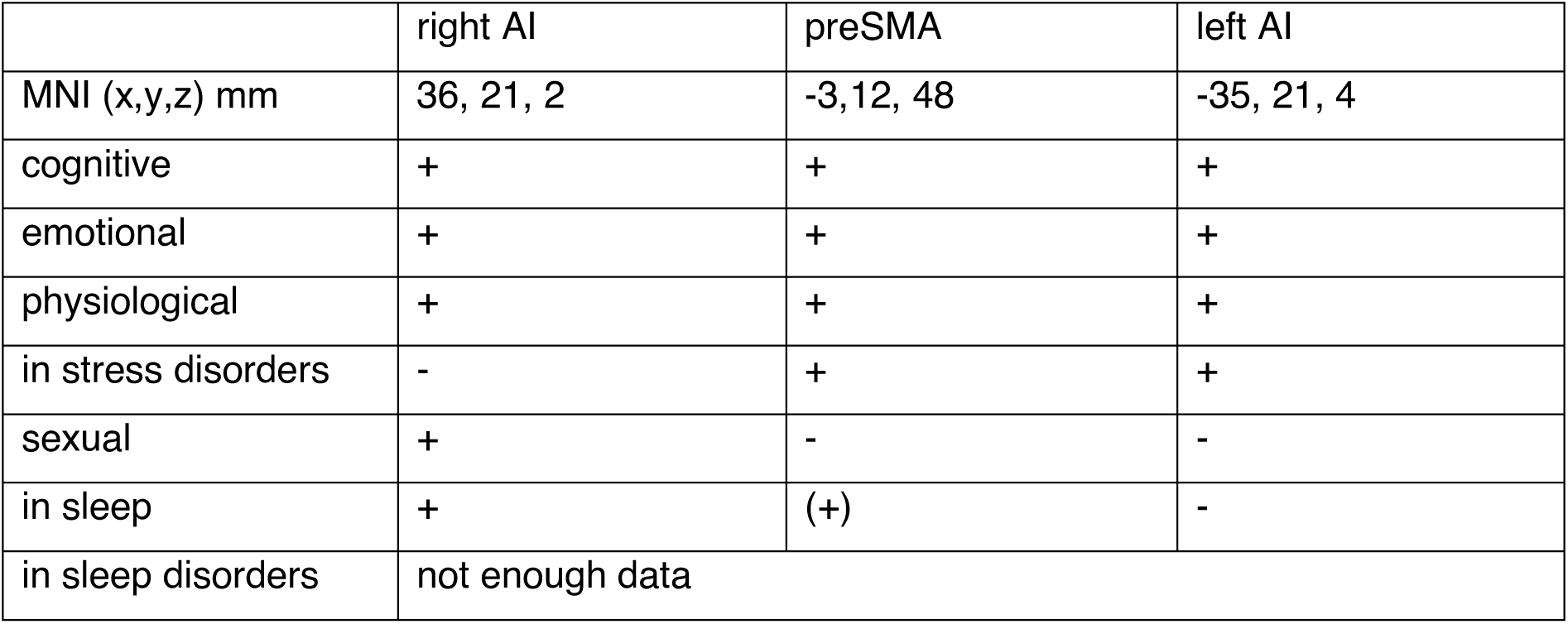
Relationship between the seven different varieties of arousal and the three nodes of the arousal cortical network, in right and left anterior insula (AI) and pre-supplementary motor areas (preSMA). Coordinates correspond to center of mass. (+) indicates the presence of significant activation and (-) the absence of significant activation (voxel-level cluster forming *p* <0.001 uncorrected; cluster level *p* <0.05 corrected for multiple comparisons using Monte-Carlo permutations). The presence of activation related to arousal in sleep in preSMA is in parenthesis because this result was not stable when varying the granularity of the semantic analysis.

The organization of data in semantic space and of data in neuronal space share some interesting similarities. The three varieties of arousal (cognitive, emotional and physiological) that are present in all the three nodes of the cortical arousal network are also close in semantic space. Arousal in sleep and arousal in stress disorders are far away from each other in semantic space, and share only the preSMA node in neuronal space. Sexual arousal, which is far from all other varieties of arousal in semantic space, differs most from the other forms of arousal in cortical space as well, since it shows up in right dorsal anterior insula only.

In the results presented above, we excluded results coming from *a priori* regions of interest. When including these results in the meta-analyses, results are overall similar (Supplementary Figure 3), but with two noticeable differences. First, there are now more studies available for the field of arousal in sleep disorders (n=16), which reveals an activation in left anterior insula. Second, a significant overlap across 6 types of arousal (all but arousal in sleep disorders) is now found in the bilateral anterior insula as well as the amygdala, covering all amygdala nuclei and some neighboring areas such as the Hippocampus or Entorhinal Cortex (Supplementary Figure 3).

#### The left and right anterior insula nodes of the cortical arousal network are both robust to the parameters of the semantic analysis and specific to arousal

We then tested whether our results were robust against changes in the parameters of the semantic analysis. The results presented above were obtained when including 1200 terms in the semantic analysis. We already showed that when varying the number of terms (or granularity) used for semantic analysis, we retrieve the same number of semantic communities with similar overall spatial organization (Supplementary Figure 1). However, increasing granularity induces two changes. First, terms which are not central to a given community but rather at the border might be reassigned to a different community. Second, with more terms in the semantic analysis, a given experimental study is likely to match more terms. Because those terms might belong to different semantic communities, a given experimental study might be assigned to more semantic communities when the granularity of the analysis increases (Supplementary Figure 1, last column). To examine these effects, we repeated the analysis described above and varied the number of terms used (1200, 1500, 2000, or 2285). We then computed in which regions there was an overlap of brain activation across four or five types of arousal, across all semantic granularity levels (Figure 3A, Supplementary Figure 4). We found that the association between left and right insula with different types of arousal was highly consistent across different granularity levels of the semantic analysis (Figure 3B). In preSMA, there was a consistent overlap of cognitive, emotional and physiological arousal as well as of arousal in stress disorders; however, the link between arousal in sleep and preSMA was not stable (Table 1).

**Figure 3.**
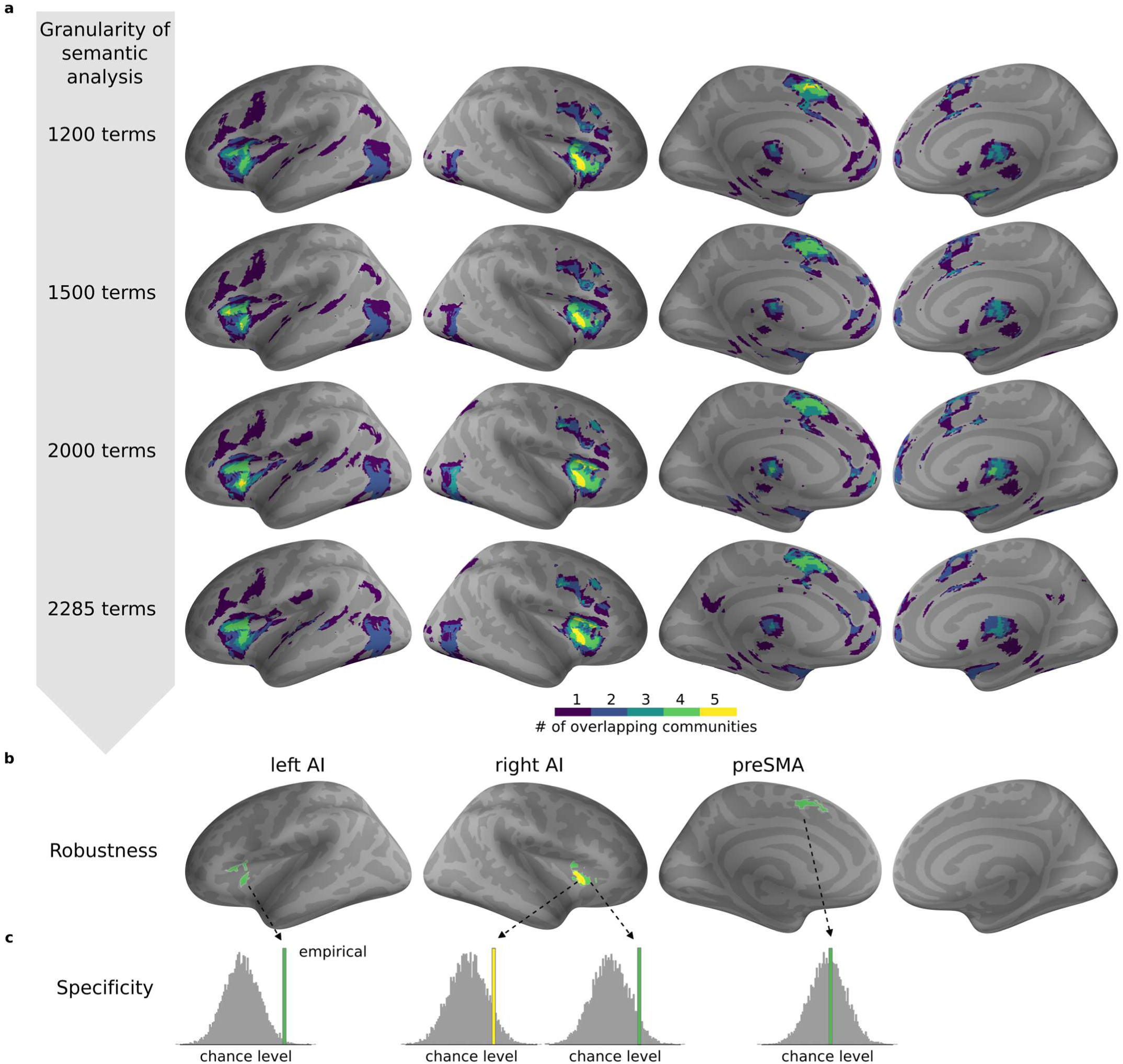
Robustness and specificity of the cortical arousal network. A. Overlap of the different communities (as last row in Figure 3) but for different levels of granularity of semantic analysis. B. Surface representation of the network robust to parameters of semantic analysis. The green color denotes areas where we found an overlap of four, and yellow an overlap of five, semantic communities across all levels of granularity of the semantic analysis. C. Specificity of each node of the robust network to arousal. For each node: left AI, right AI and preSMA, we tested how likely an activation in the region was in the non-arousal literature (gray histogram, chance level distribution) as compared to the arousal literature (colored bar). *Abbreviations*: AI (anterior insula), preSMA (pre-supplementary motor area).

Finally, we tested whether the activation of the three nodes of the arousal cortical network was *specifically* associated with arousal. Indeed, both the anterior insula^18–20^ and the preSMA^21^ (often labelled dorsal anterior cingulate cortex or dorso-medial prefrontal cortex dACC/dmPFC) are notoriously often activated in *any* paradigm, including in paradigms not particularly designed to target arousal. We thus assessed whether a significant activation cluster could be obtained by chance in those three nodes. We defined chance level by activation likelihood in studies not using the term arousal. For each node, we computed a maximal statistic (see Materials & Methods) across the 415 arousal related studies. We then computed the same statistic in 415 studies not related to arousal randomly drawn from the NeuroQuery database, and repeated the process 10.000 times to obtain a chance level distribution. Because this analysis includes studies using a region of interest approach, results might be slightly distorted, and p-values are only indicative. Still, there is a marked difference between preSMA, where the arousal-related results correspond to what could be expected when selecting studies randomly (Figure 3C), and the left and right anterior insula arousal-related nodes, which appear to be significantly (left AI, Monte-Carlo p=0.0065) or marginally (right AI(5), p=0.082, right AI(4), p=0.05971) specific to arousal.

We conclude that the most robust and specific association with arousal is found in the left and right anterior insula. The arousal-related nodes in the anterior insula were anatomically symmetrical (Figure 4 A, B), although they correspond to the convergence of different types of arousal (Table 1). We further characterized the insular arousal hubs. Anatomically, the insular arousal hubs were confined to the anterior short gyrus^22^ (Figure 4C), in a dysgranular dorsal anterior territory, centered in area Id7 and extending dorsally in Id6 as well as marginally ventrally in Id8^23–25^ (Figure 4B). In terms of resting state networks, the dorsal insula arousal hubs belong to the Ventral Attention Network^26^ (Figure 4D). They overlap more specifically with the typical seed used for the cingulo-opercular network^27^, but are more dorsal than the seed region of the homeostatic salience network^28^. The dorsal location of the insula arousal hubs is confirmed when comparing our results with the tripartite organization (dorsal cognitive, ventral emotional and posterior sensorimotor) of the insula based on both resting state functional connectivity^29^ and meta-analytic task-related activations^30^ (Figure 4E). Finally, the arousal hubs in dorsal anterior insula are included in a larger region related to predominantly parasympathetic regulation of the autonomic nervous system during tasks, and are more dorsal than the predominantly sympathetic regulation^31^ (Figure 4F).

**Figure 4.**
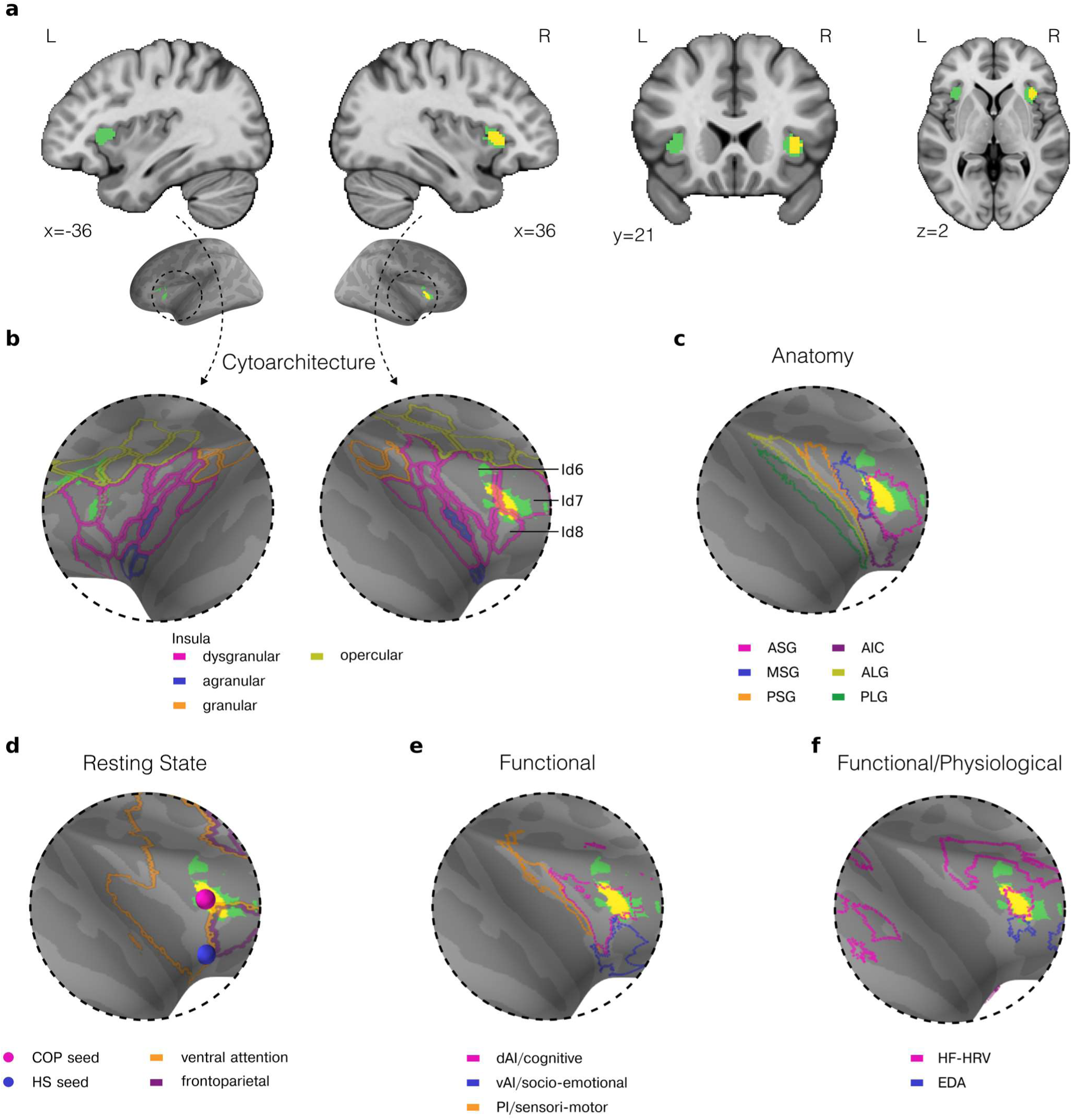
Dysgranular area 7 in dorsal anterior insula is a key hub for arousal. A. Volume representation of the robust and specific association with arousal in left and right anterior insula. B. Surface representation of the right and left anterior insula with description of the cytoarchitecture defined in the Julich-Brain^23–25^ C. Location of the most robust node in the right anterior insular cortex, with respect to the anatomical parcellations of Faillenot et al.^22^, D. resting state connectivity networks as defined in Yeo et al.^26^, and seeds used to define cingulo-opercular (COP) network in Dosenbach et al.^27^, and the homeostatic salience (HS) network in Seeley et al.^28^, as well as E. functional parcellation from Deen et al.^29^ with functional description from Kurth et al.^30^, and F. contours of sympathetic (EDA) and parasympathetic (HRV) task-evoked changes as described in the meta-analysis of Beissner et al.^31^. *Abbreviations*: Id (dysgranular insula); ASG (Anterior Short Gyrus), MSG (Middle Short Gyrus), PSG (Posterior Short Gyrus), AIC (Anterior Inferior Cortex), ALG (Anterior Long Gyrus), PLG (Posterior Long Gyrus), dAI (dorsal Anterior Insula), PI (Posterior Insula), vAI (ventral Anterior Insula), COP (cingulo-opercular), HS (homeostatic salience), HF-HRV (high frequency heart rate variability), EDA (electrodermal activity).

## Discussion

We identify in a purely data-driven manner seven varieties of arousal: cognitive, emotional, physiological, and sexual arousal, as well as arousal in sleep, in sleep disorders and in stress disorders. The seven varieties of arousal differ not only in semantic space but also in both physiological measures and specific cortical correlates. Yet, except for arousal in sleep disorders where data are scarce, all varieties of arousal converge, albeit with different weights, onto three nodes of a cortical arousal network composed of left and right anterior insula and preSMA. The amygdala is likely to be part of this network, but because it is most often isolated through a region of interest analysis likely to distort meta-analyses^16,17^, we limit our conclusions to cortical regions. Our results show that even if the different varieties of arousal refer to markedly distinct domains, there is enough commonalities in the underlying phenomenon for neuronal data to converge on the same cortical arousal network. Among the nodes of the arousal network, the ones located in left and right dorsal anterior insula were not only robust against changes in the granularity of the semantic analysis, but they were also specific to arousal. These hubs of convergence of different arousal varieties were located in the anterior short gyrus of the anterior insula, specifically in dysgranular area Id7. While cognitive, emotional and physiological arousal were present in both left and right dorsal anterior insula Id7, sexual arousal and arousal in sleep were right lateralized, and arousal in stress disorders was left lateralized. Our results thus indicate that a common network underlies the notion of arousal across very different domains, and reveal the central position of dysgranular insula area Id7 in domain-general arousal.

### Situating the cortical arousal network in the brain network literature

Arousal refers to a brain-body state and is known to involve ascending neuromodulatory systems. The three nodes of the cortical arousal network we identify here are functionally connected to ascending neuromodulatory systems^32^, but represent only a subset of a much larger ensemble that also comprises fronto-polar and fronto-orbital regions^32^. The cortical arousal network also overlaps with regions co-varying with bodily measures of autonomic activity during tasks^31^, in line with the notion that arousal reflects a mobilization of brain and body resources. More precisely, the preSMA node co-varies with measures of sympathetic activity and the dorsal insula nodes co-vary with high-frequency heart rate variability, a measure of parasympathetic activity. However the cortical autonomic network^31^ is much larger than the arousal network we reveal here. The cortical arousal network is thus related to, but not confounded with, targets of ascending neuromodulatory systems, and regions involved in autonomic functions.

The nodes of the arousal network are part of a large resting-state network often referred to as the ventral attention network^26^ or salience network^33^. Some authors further subdivide this network in two, the cingulo-opercular network^27^, and the homeostatic salience network^28,34^. Within the anterior insula, the homeostatic salience network^28,34,35^ is typically more ventral than the arousal hub we identified. Our results appear better aligned with the cingulo-opercular network, which is more dorsal in the anterior insula. The cingulo-opercular network has been related to task set maintenance^27^, and to a state of tonic alertness, or “self-initiated (rather than externally driven) preparedness to process and to respond”^36^. While the anatomical distinction between homeostatic salience and cingulo-opercular networks in the anterior insula is still debated^33^, both networks are considered to play a role in regulating the integration or segregation of other brain networks^37^, a role also ascribed to arousal^38,39^.

The arousal network is composed of brain regions known to be very often activated irrespective of the specific nature of the task^18,20,21^. Arousal could be the underlying domain-general variable explaining those activations. Still, this interpretation has to be nuanced. Indeed, we find that the arousal node in preSMA is not specific to arousal. Such a finding might be interpreted in two different ways: one is that arousal, construed as a mobilization of brain and body resources, is an underlying factor in most if not all tasks, even when arousal is not an explicit target of the experimental study. In this view, arousal would be a fundamental dimension to consider when interpreting any task-related activation. Alternatively, it could be that the association between preSMA and arousal in our results is driven by another factor common to both arousal and non-arousal studies. The present results do not allow to disambiguate between these two interpretations in preSMA. However, in the case of the dorsal anterior insula nodes of the arousal network, the interpretation is much more straightforward, since the dorsal Anterior Insula nodes were specific to studies explicitly mentioning arousal.

### The robust and specific nodes of the arousal network: Id7 in the dorsal anterior insula

Dysgranular area Id 7 in the left and right dorsal anterior insula appears as a key hub for arousal for three reasons. First, it reflects the convergence of the highest number of varieties of arousal (cognitive, emotional, physiological, sexual, sleep-related in the right hemisphere; cognitive, emotional, physiological, in stress disorders in the left hemisphere). Second, this convergence was robust to changes in the granularity of the semantic analysis. Last but not least, activity in this region is specific to arousal – in other words, this region is more likely to be activated when arousal is mentioned in the article than when arousal is not mentioned. Area Id 7 has only been recently anatomically identified in humans^22^ and functional studies are scarce, but it constitutes a promising novel seed for large-scale brain analysis^35,40^ to refine the definition and mechanisms of domain-general arousal.

Anatomo-functional specification within the insula, while recognized as important^18,41^, has often been limited in the experimental literature in humans to a rough partition into three regions, typically posterior, dorsal anterior, and ventral anterior^29,42^. Recently however, anatomical studies in humans^24^ and non-human primates^43^ revealed a much finer grained organization. Anatomical specification is all the more important that the anterior insula is disproportionately enlarged in humans compared to other species^44^. To the best of our knowledge, whether the human Id7 has a homologue in non-human species is not yet known. Functionally, the electrical stimulation of the posterior insula in human patients can elicit various physiological changes and/or somato-visceral sensations^45,46^. In contrast, the stimulation of the anterior part of the insula^46^, and specifically of the right anterior insula area Id7^47^, does not elicit any noticeable response in human patients at rest. This suggests that Id7 is neither a primary viscero-sensory nor viscero-motor node. However Id7 becomes active in a simple visual oddball task, with infrequent non-target visual scenes eliciting large neuronal responses^47^, suggesting this area is able to keep track of task structure to detect deviant stimuli.

### Hemispheric lateralization of Id7 in dorsal anterior insula

We observe some hemispheric specialization for arousal in dorsal anterior insula: while cognitive, emotional and physiological arousal are bilateral, arousal in stress disorders appears in left Id7 and sexual and sleep-related arousal in right Id7. While a functional asymmetry is often observed during tasks between the left and right dorsal anterior insula (see, e.g., Macey et al.^48^), there seems to be no marked lateralization in either cytoarchitectonic organization^49^ nor cortical resting-state functional connectivity^50^. However, both functional connectivity^51^ and tractography^52^ data suggest a lateralization of the connectivity between anterior insular cortices and basal ganglia.

Arousal in stress disorders is the only variety of arousal not present in the right insula. The fMRI studies of arousal in stress disorders included in the meta-analysis cover quite a broad range of disorders, including anxiety and depression but also bipolar disorder, borderline personality disorder, obsessive compulsive disorder, etc. Despite the heterogeneity of the disorders, all those studies converge in the preSMA and left insula, which shows that they have enough in common to activate two nodes of the cortical arousal network. However, they fail at providing any evidence, even at subthreshold level, for an activation in the right anterior insula. One finding that seems recurrent in the stress disorder literature is that there is a partial decoupling between subjective reports and peripheral measures of arousal in patients, with for instance limited objective signs of autonomic hyperarousal at rest in chronic anxiety patients^53^, and only a weak association between subjective reports of sleep disturbance and objective polysomnographic measures of sleep in post-traumatic stress disorder^54^. A better understanding of the functional lateralization of arousal in dorsal anterior insula might thus require refining how arousal is measured, objectively or subjectively, with potential clinical implications.

### Conclusion: Domain-general arousal reconciles the global state view with the multi-dimensional construct view

We provide here evidence for the existence of domain-general arousal at the cortical level and highlight dysgranular area 7 as a critical hub that may act by regulating the integration and segregation of other brain networks. Domain-general arousal is compatible with, but not synonymous with general arousal construed as a global state^1,6^. Indeed, general arousal is often presented as a process that globally affects brain processing and behavior at a given moment in time. What we show here is that arousal in situations as different as a cognitive task, an emotional context, the transition to wakefulness, or sexual behavior, occurring at different moments, are associated with a common neural network. Because we analyzed only cortical correlates of arousal, our results are mute on the involvement of subcortical pathways specific to some varieties of arousal, but our results are compatible with the notion that there are cortical differences between varieties of arousal^7^. Domain-general arousal should thus become a central component of any taxonomy of arousal since it can reconcile the seemingly opposing concepts of general and multi-dimensional arousal: the activation of the domain-general arousal cortical network might be obtained via different neuroanatomical routes in different experimental situations – a view compatible with the multi-dimensional construct – and lead to a global reconfiguration of brain networks – a view compatible with the global state of general arousal.

## Materials & Methods

### Semantic analysis

#### Data collection

We collected the 122 590 records returned by the exact query “arousal” searched in all available fields from Web of Science (46 924, search on May 5th, 2023) and Pubmed (53 211, search on April 10th, 2023 using Pubmed Entrez Direct command line utility (https://ftp.ncbi.nlm.nih.gov/entrez/entrezdirect/)). Note that by default the Pubmed API extends queries by semantic similarity (i.e., the query ‘arousal’ extends to ‘wakefulness’), but here we enforced the exact query mode. We excluded records that were missing data in any of the following fields: ‘abstract’, ‘title’, ‘pubmed id’, ‘language’, “document type” or ‘publication date’. We then selected records corresponding to original research articles written in English. Finally, we removed duplicates based on PubMed ID. This resulted in a dataset composed of 49 525 unique records.

#### Semantic network

To explore the semantic landscape of the arousal literature we used the bunkatech library^45^ designed for Natural Language Processing applications. The bunkatech term extractor uses Textacy (https://textacy.readthedocs.io/en/latest/), a library built on top of Spacy Deep Learning Language model (https://spacy.io/models/en). The term extractor was parameterized to extract the most frequent terms composed of 2 or 3 words, to include nouns, proper nouns, and adjectives, and we recovered the 3500 terms most commonly occurring in the dataset.

A number of terms were excluded from further analysis (see Supplementary Table 1 for exclusion criteria) because they were too general (i.e., ‘control group’ or ‘significant results’). We also removed all brain related terms like ‘brain imaging’, ‘prefrontal cortex’ etc., by matching with NeuroNames^46^ dictionary and further manual exclusion. Term exclusion was performed manually by authors MS and CTB separately, disagreements were discussed until resolved, to obtain a final list of 2285 terms, sorted by occurrence. The final list of the 2285 terms retained is available on OSF.

The following steps were completed for variable number of terms (300, 600, 900, 1200, 1500, 2000, 2285). We first created a co-occurrence matrix of the terms, with co-occurrence being defined within the same entry in the dataset (i.e., terms appearing in the same abstract). We then computed the semantic similarity (also called semantic distance) between the terms using cosine similarity (computed on the co-occurrence), and retained the 15 closest neighbors of each term. The result is an edge table consisting of nodes (terms) and edges (cosine distance) between the nodes. We embedded the edge table into 700 dimensions using Node2Vec. The nodes were then clustered using K-means. The ‘number of clusters’ parameter was optimized using the elbow method. Finally, the dimensionality was reduced with UMAP (Uniform Manifold Approximation and Projection for Dimension Reduction)^55^ for visualization.

#### Article-cluster assignment

We associated scientific articles with one of the seven semantic clusters based on the terms detected in the abstracts. If all the terms detected in the abstract belonged to one semantic cluster, the corresponding article was associated with this cluster. If an abstract was indexed with two terms belonging to two different semantic clusters, i.e. ‘heart rate’ and ‘stress disorder’, the corresponding article was associated with two clusters, in this example ‘physiology’ and ‘stress disorders’.

### Semantic prevalence of physiological arousal measures

To calculate the prevalence of the physiological measure of arousal in each cluster we searched for words from a dictionary of related terms we created (see Supplementary Table 3). We matched the words using simple python string matching by checking whether the term in the dictionary is present in the dataset item. The final result is the proportion of articles in which the relevant terms were found present over all the articles in the semantic cluster.

### fMRI meta-analysis-Activation Likelihood Estimation

#### Data Selection

For the neuroimaging meta-analysis we used the open-access neuroimaging peak-coordinate dataset available from NeuroQuery^34^. We selected eligible studies from our ‘arousal’ literature dataset by matching pubmed ID with the Neuroquery dataset. We included only original studies reporting coordinates of brain activations, i.e., from the 429 studies available in the NeuroQuery database, we included EEG, MEG and fMRI studies but excluded 7 morphometry studies, 1 DTI study, 5 meta-analyses and 1 article that did not report any coordinates. Because NeuroQuery relies on automatic extraction of coordinates from tables, we further verified which of those 415 studies reported activation coordinates resulting of a region of interest analysis (ROI) in a table. This is an important step to reduce the bias for oversampling a given region (e.g., amygdala in emotion studies). For each paper we assessed Abstract, Materials and Methods, Tables, and when necessary Results to select studies where coordinates in tables corresponded to results of whole brain analysis, or used functionally defined ROI. We identified 187 articles where tables reported coordinates of results based on a priori defined ROIs, leaving a dataset of 228 whole-brain neuroimaging studies. Most of those studies contained at least one of the terms of the semantic network and could thus be associated to at least one semantic cluster. The exact number of studies that could be associated with semantic clusters depends on the number of terms used in the semantic analysis, for instance with 1200 terms in the semantic network 221 neuroimaging studies contained at least one term used in the semantic network and could thus be assigned to a least one semantic cluster.

#### Sample size

We used the online application Elicit to retrieve information on sample sizes for the neuroimaging papers. For the articles for which the language model was not successful we noted the sample size by hand.

#### ALE meta-analysis

Coordinate-based quantitative meta-analyses of neuroimaging results were performed using NiMARE v0.0.11^47,48^ leveraging its integration with the NeuroQuery database^34^. Activation Likelihood Estimation (ALE) is based on computing consistent activation foci by modeling the probability distribution of activation at given coordinates against the null distribution of summary-statistic values and their expected frequencies under the assumption of random spatial associations between studies, via a weighted convolution^36^. We ran a separate ALE meta-analysis for each semantic field, with the following parameters: fixed full-width half-max of 10 for Gaussian kernel, cluster forming uncorrected significance level of 0.001, and Cluster-level inference with correction for multiple comparisons using family wise error correction with Monte-Carlo permutations to identify brain areas consistently activated across articles of interest. The results of the meta-analyses were represented in MNI152 coordinates, that were then transformed into freesurfer (http://surfer.nmr.mgh.harvard.edu/) inflated surface representation using pysurfer (v0.11.0) (https://pysurfer.github.io/index.html) vol2surf function for visualization. The maps were thresholded for significance at one-sided Monte-Carlo alpha level = 0.05.

#### Convergence between research fields

To identify structures common across the results of meta-analyses in each cluster, we overlapped the significance maps across semantic clusters and identified brain regions where the highest convergence between semantic fields was observed. To assess the robustness of resulting overlap foci against variations in semantic analysis parameters, we ran the fMRI meta-analysis following a semantic analysis with a variable number of terms: 1200, 1500, 2000, or 2285. We created an overlap map for each number of terms. To determine the probability that an overlap of 4 (or 5) semantic communities occurs at a given location irrespective of the granularity of the semantic analysis (robustness maps), we combined the overlap maps obtained from different number of terms in the semantic analysis. In practice, we created a mask for 4 (or 5) communities in overlap at each granularity level, and created a conjunction across granularity levels.

We report MNI152 coordinates of the center of mass (computed using ndimage.center_of_mass function from SciPy library (v1.11.2) ^56^). For visualization we projected resulting maximal overlap foci to volume using the freesurfer surf2vol function and reported the MNI152 coordinates. The identified foci were visually compared with existing parcellations in surface. Surface projections were available for Julich-Brain^23–25^ (v2.9 https://search.kg.ebrains.eu/instances/61c09a8a-bbfe-49eb-ab45-354b05f9a600) and Yeo et al^26^. 7 network projection (included in the freesurfer package). The parcellations from Deen et al.^29^, and Faillenot et al.^22^ (www.brain-development.org/brain-atlases/adult-brain-atlases/), were available in volume, we projected them to surface using the freesurfer vol2surf function.

#### Maximal statistic and null distribution

We selected cluster mass as the maximal statistic. Cluster mass refers to the sum of summary statistic (in our case the z-transformed t-test) across all voxels in the cluster. This statistic is often used for cluster inference – the cluster mass null distribution can be generated with permutation tests, and then the tested cluster mass significance is calculated based on that null distribution. In practice, we created overlap regions of interest (ROIs) using the *robustness* maps – for each area where we find a consistent overlap of four (or five) semantic communities. For each overlap ROI, we computed the maximal statistic for all 415 arousal related articles (including articles with an a priori ROI approach). We then established the null distribution by repeatedly selecting sets of 415 articles not explicitly mentioning arousal and computed cluster mass within the overlap ROI. The only criteria for an article to be included in the chance level estimation was that it should not contain the word arousal; the article could explicitly mention emotion, or sleep, or pupil diameter etc…and could include an a priori ROI approach. We repeated this ALE analysis 10.000 times to generate the null distribution of activation likelihood when arousal is not mentioned. We then compared the value obtained in overlap ROIs for the 415 arousal-related articles to the null distribution to obtain a Monte-Carlo p.

## Data Availability

The data and code to reproduce results in the manuscript will be published online upon acceptance in a peer-reviewed journal.

## Acknowledgements

This work was supported by FrontCog grant number ANR-17-EURE-0017 and a fellowship of the Canadian Institute For Advance Research (CIFAR) program in Brain, Mind and Consciousness to C.T.-B. C.D was supported by CNRS-University of Tokyo grant to Hugo Mercier (“Science communication for and by citizens”). We thank Karen Quigley for providing guidance at an early phase of the project, Julien Karadayi for sharing his expertise in graph analysis, Andrei Mogoutov for advice on natural language analysis, Thomas Andrillon, Bence Farkas, Julie Grèzes, Marie Loescher, Alizée Lopez-Persem, and Stefano Palminteri for useful comments on the initial version of the manuscript.

A CC-BY public copyright license has been applied by the authors to the present document and will be applied to all subsequent versions up to the Author Accepted Manuscript arising from this submission, in accordance with the ANR grant’s open access condition.

## Authors contributions

MS had the initial idea of using semantic analysis; MS & CTB designed the project; MS and CdD ran the semantic analysis, MS and CTB the brain imaging meta-analysis. MS and CTB wrote the first version of the article, all authors contributed to the final version of article.

