## Supplementary Materials for "Evidence for domain-general arousal from semantic and neuroimaging meta-analyses reconciles opposing views on arousal"

### Different varieties of arousal converge on a shared cortical network

Supplementary Figure 1 – Semantic network for different granularities
Supplementary Figure 2 – Meta-analyses as in Main Fig 2, unthresholded
Supplementary Figure 3 – Meta-analyses as in Main Fig 2, region of interest included
Supplementary Figure 4 – Neuroimaging meta-analyses for 1500, 2000, 2288 terms in semantic analysis

Supplementary Table 1 – Guidelines for term inclusion
Supplementary Table 2 – Terms with largest inside degree centrality
Supplementary Table 3 – Dictionary of physiological measures

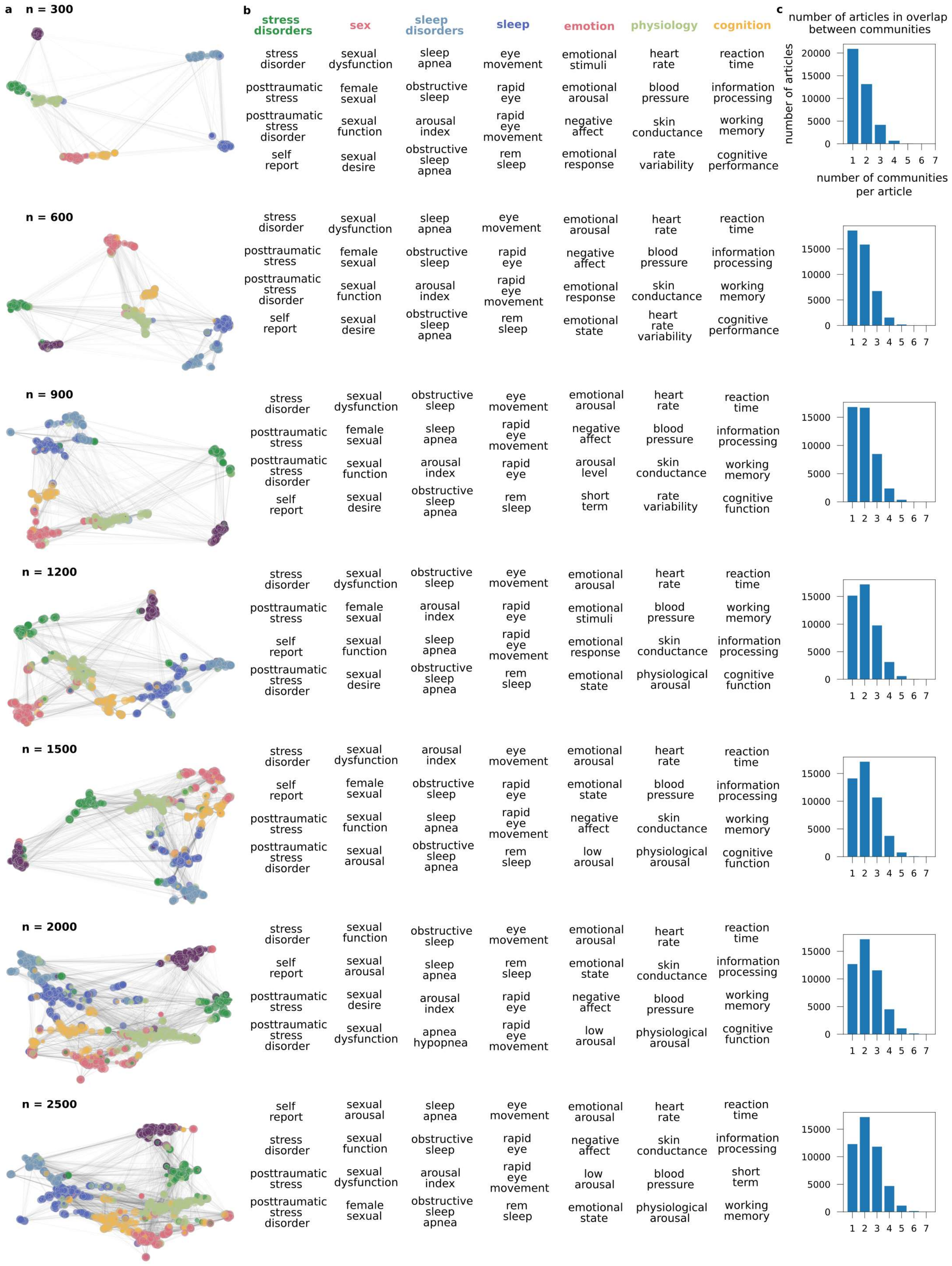

a
n = 300
n = 600
n = 900
n = 1200
n = 1500
n = 2000
n = 2500
b
stress disorders
sex
sleep disorders
sleep
emotion
physiology
cognition
stress disorder
sexual dysfunction
sleep apnea
eye movement
emotional stimuli
heart rate
reaction time
posttraumatic stress
female sexual
obstructive sleep
rapid eye
emotional arousal
blood pressure
information processing
posttraumatic stress disorder
sexual function
arousal index
rapid eye movement
negative affect
skin conductance
working memory
self report
sexual desire
obstructive sleep apnea
rem sleep
emotional response
rate variability
cognitive performance
stress disorder
sexual dysfunction
sleep apnea
eye movement
emotional arousal
heart rate
reaction time
posttraumatic stress
female sexual
obstructive sleep
rapid eye
negative affect
blood pressure
information processing
posttraumatic stress disorder
sexual function
arousal index
rapid eye movement
emotional response
skin conductance
working memory
self report
sexual desire
obstructive sleep apnea
rem sleep
emotional state
heart rate variability
cognitive performance
stress disorder
sexual dysfunction
obstructive sleep
eye movement
emotional arousal
heart rate
reaction time
posttraumatic stress
female sexual
sleep apnea
rapid eye movement
negative affect
blood pressure
information processing
posttraumatic stress disorder
sexual function
arousal index
rapid eye
arousal level
skin conductance
working memory
self report
sexual desire
obstructive sleep apnea
rem sleep
short term
rate variability
cognitive function
stress disorder
sexual dysfunction
obstructive sleep
eye movement
emotional arousal
heart rate
reaction time
posttraumatic stress
female sexual
arousal index
rapid eye
emotional stimuli
blood pressure
working memory
self report
sexual function
sleep apnea
rapid eye movement
emotional response
skin conductance
information processing
posttraumatic stress disorder
sexual desire
obstructive sleep apnea
rem sleep
emotional state
physiological arousal
cognitive function
stress disorder
sexual dysfunction
arousal index
eye movement
emotional arousal
heart rate
reaction time
self report
female sexual
obstructive sleep
rapid eye
emotional state
blood pressure
information processing
posttraumatic stress
sexual function
sleep apnea
rapid eye movement
negative affect
skin conductance
working memory
posttraumatic stress disorder
sexual arousal
obstructive sleep apnea
rem sleep
low arousal
physiological arousal
cognitive function
stress disorder
sexual function
obstructive sleep
eye movement
emotional arousal
heart rate
reaction time
self report
sexual arousal
sleep apnea
rem sleep
emotional state
skin conductance
information processing
posttraumatic stress
sexual desire
arousal index
rapid eye
negative affect
blood pressure
working memory
posttraumatic stress disorder
sexual dysfunction
apnea hypopnea
rapid eye movement
low arousal
physiological arousal
cognitive function
self report
sexual arousal
sleep apnea
eye movement
emotional arousal
heart rate
reaction time
stress disorder
sexual function
obstructive sleep
rapid eye
negative affect
skin conductance
information processing
posttraumatic stress
sexual dysfunction
arousal index
rapid eye movement
low arousal
blood pressure
short term
posttraumatic stress disorder
female sexual
obstructive sleep apnea
rem sleep
emotional state
physiological arousal
working memory
c
number of articles in overlap between communities
number of articles
number of communities per article
20000
15000
10000
5000
0
1 2 3 4 5 6 7

**Supplementary Figure 1. The organization of the semantic network remains stable when varying the number of terms included in the semantic analysis**. We conducted the semantic analysis on a number of terms varying from 300 to 2285. The number of clusters and their semantic contents remains stable. **A.** Semantic network and clusters. **B.** The four most occurring terms in each cluster. **C.** Histograms describing the number of semantic clusters each article is associated with.

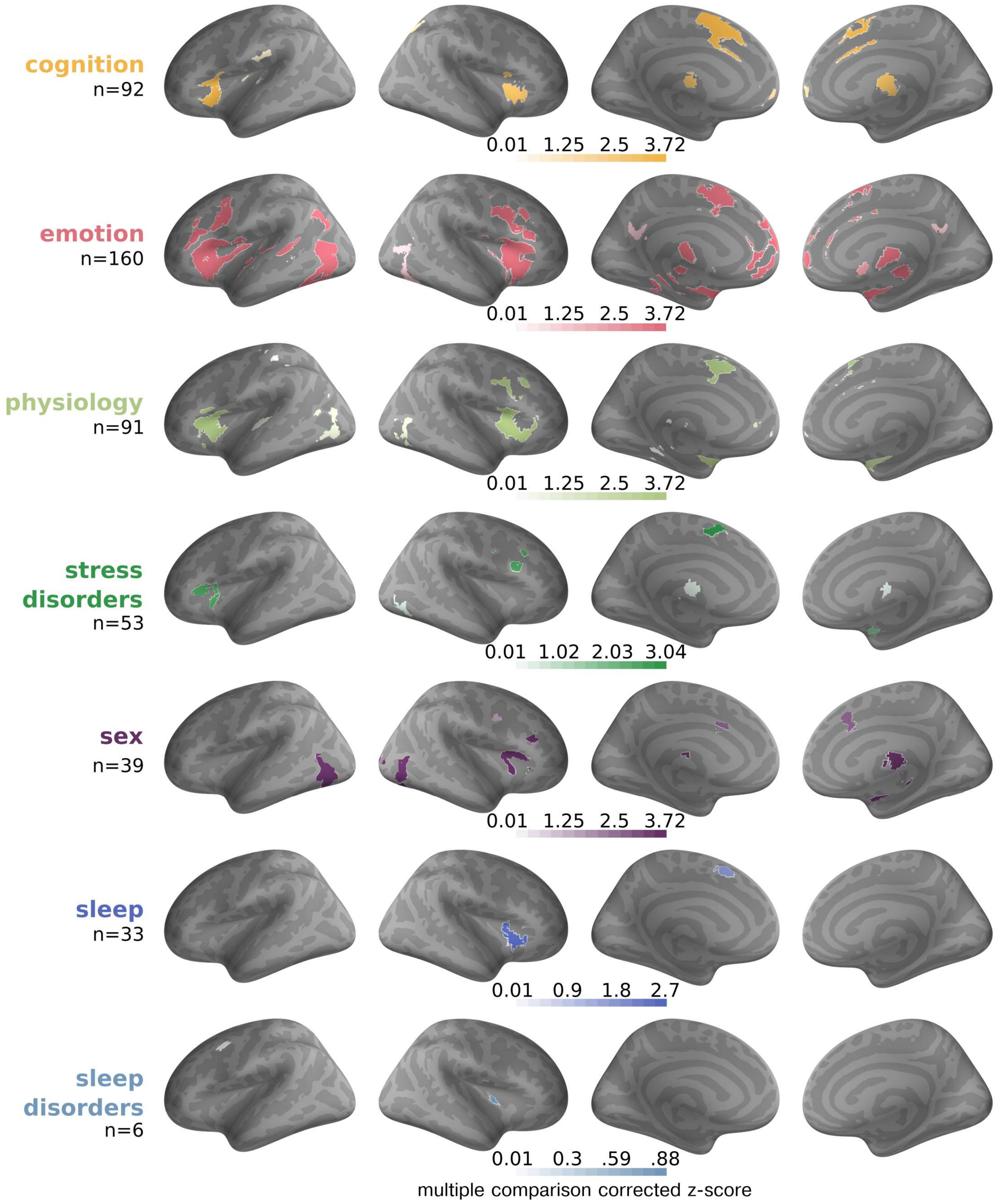


**Supplementary Figure 2. Unthresholded results of meta-analyses** (z-score, corrected for multiple comparisons) of the neuroimaging meta-analyses reported in Figure 2. Unthresholded data do not reveal more overlap in the cortical arousal network.

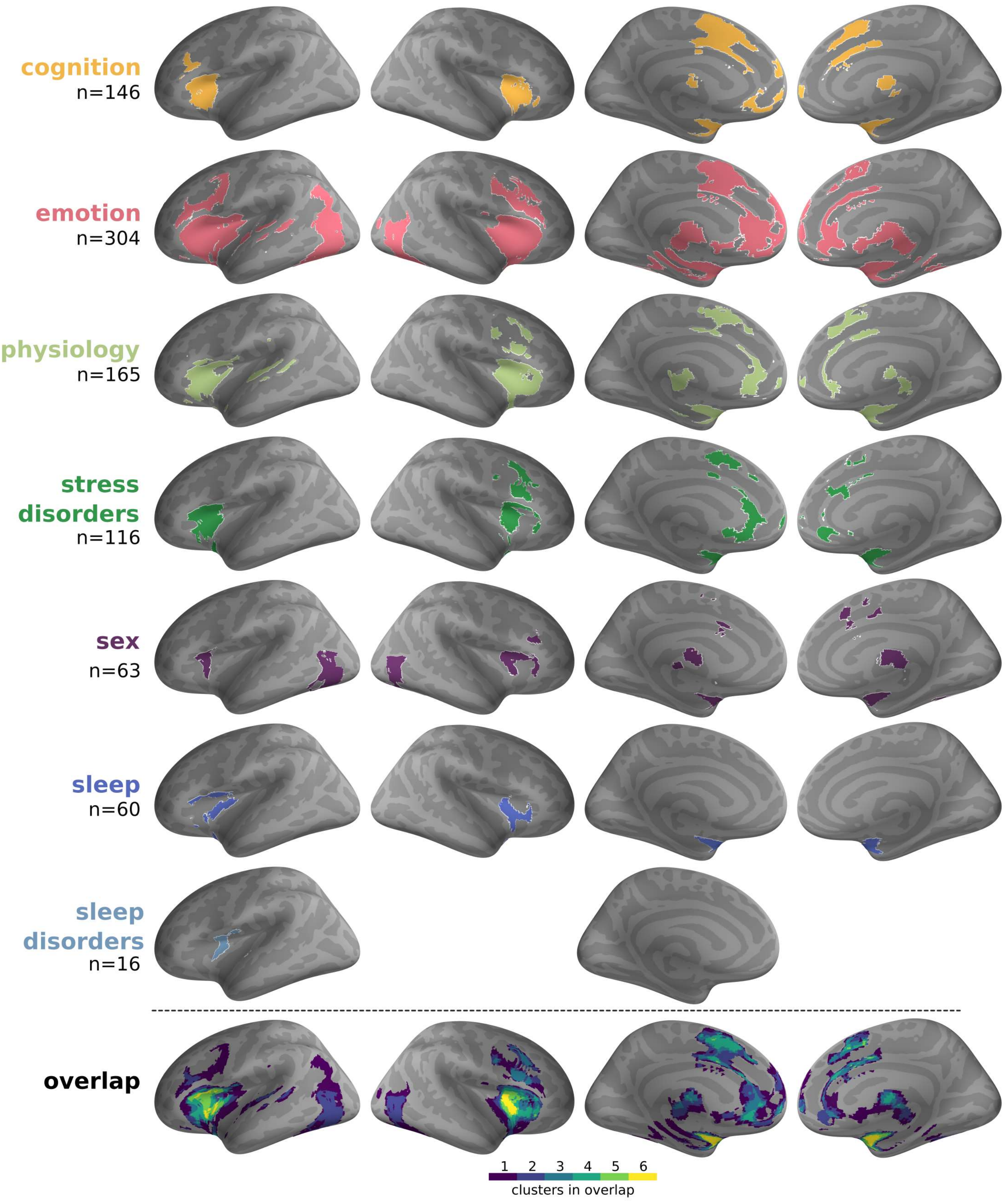


**Supplementary Figure 3. Results of the neuroimaging meta-analysis in the network of 1200 terms with articles that used a priori region of interest analysis.** Including fMRI studies that used a priori region of interest analysis in the meta-analysis maintains the overlap in the left and right anterior insula and preSMA, but reveal an additional cluster in the amygdala bilaterally.

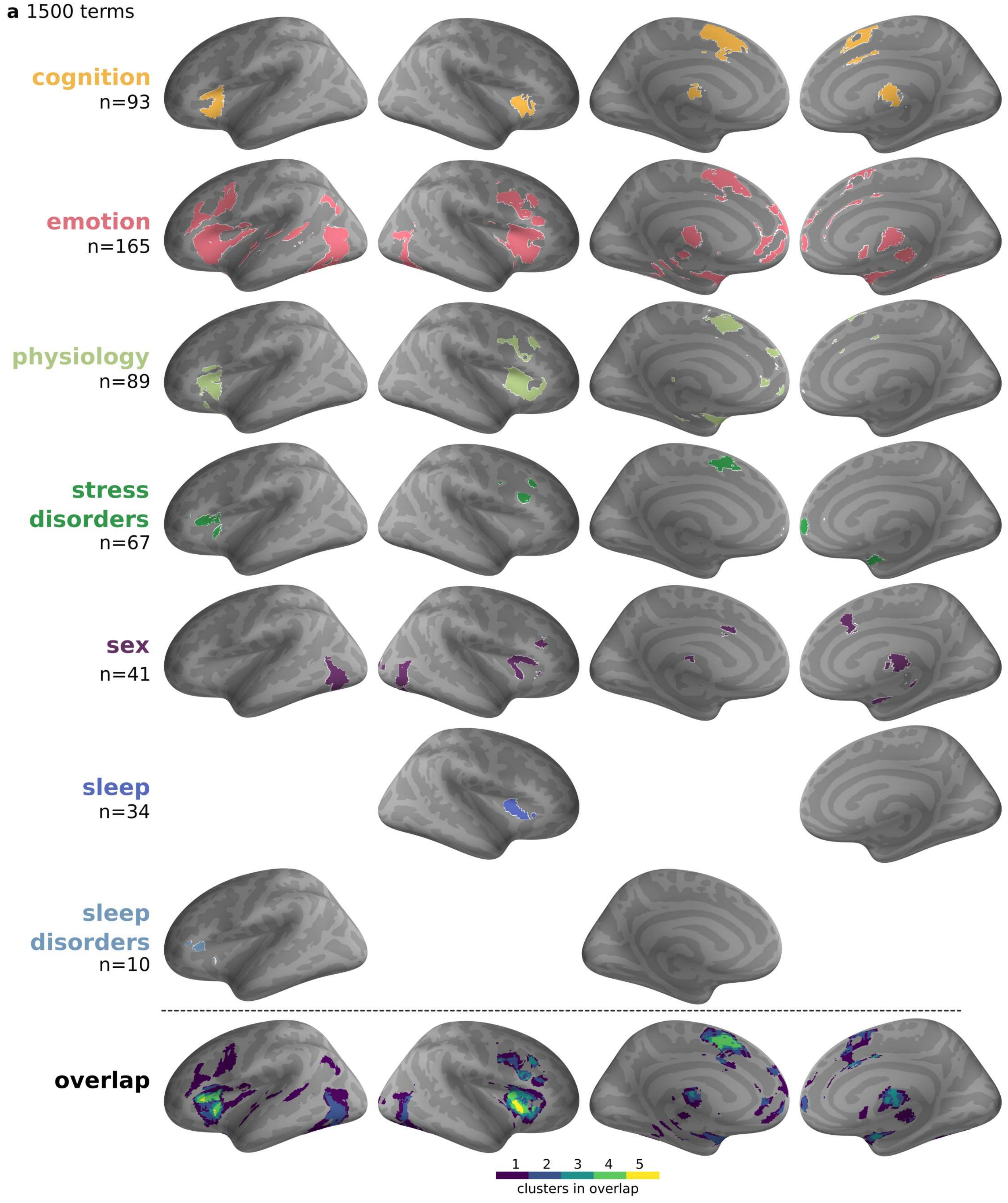

a 1500 terms
cognition
n=93
emotion
n=165
physiology
n=89
stress disorders
n=67
sex
n=41
sleep
n=34
sleep disorders
n=10
overlap
1 2 3 4 5
clusters in overlap


30

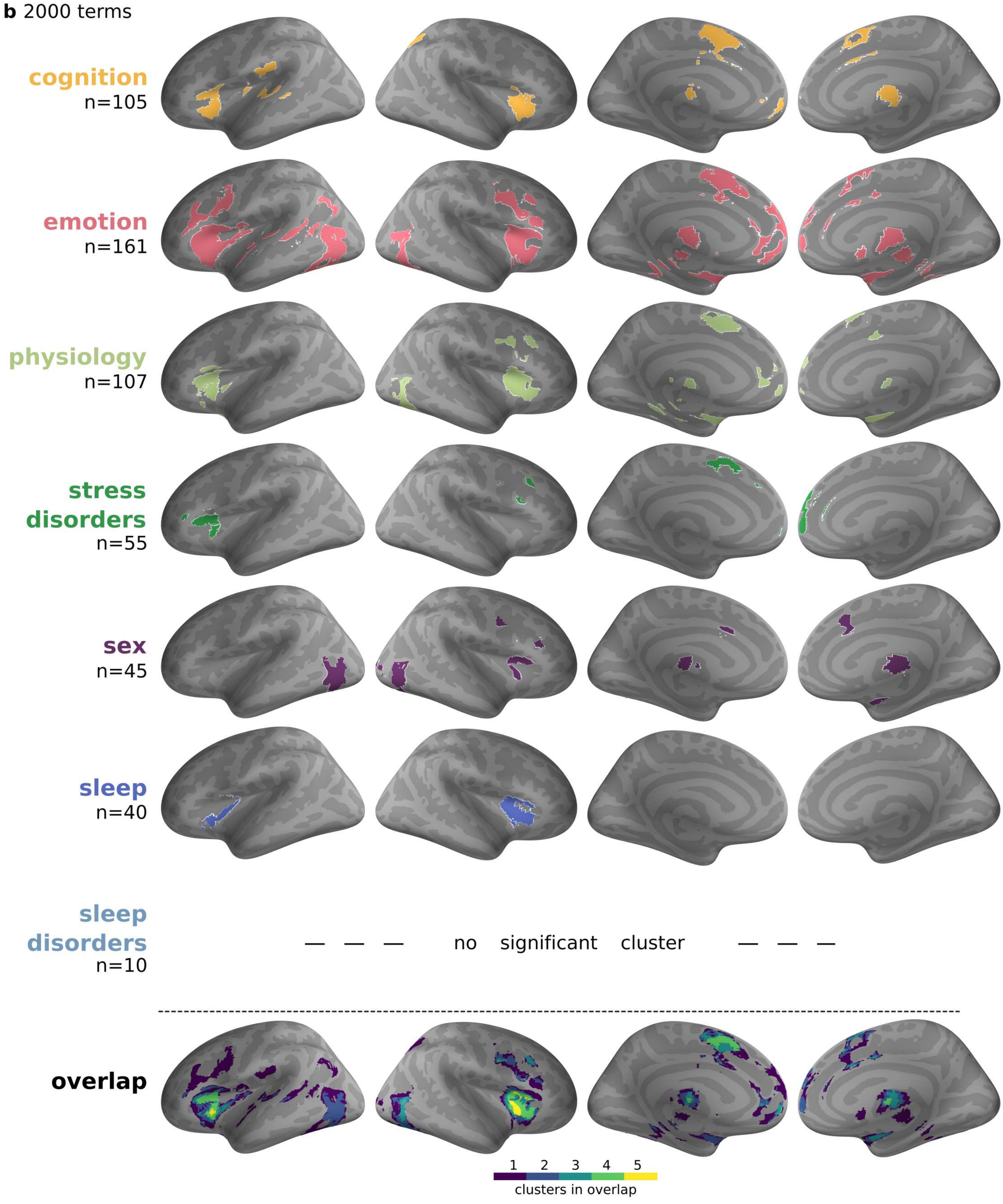
b 2000 terms
cognition
n=105
emotion
n=161
physiology
n=107
stress
disorders
n=55
sex
n=45
sleep
n=40
sleep
disorders
n=10
— — — no significant cluster — — —
overlap
1 2 3 4 5
clusters in overlap

31

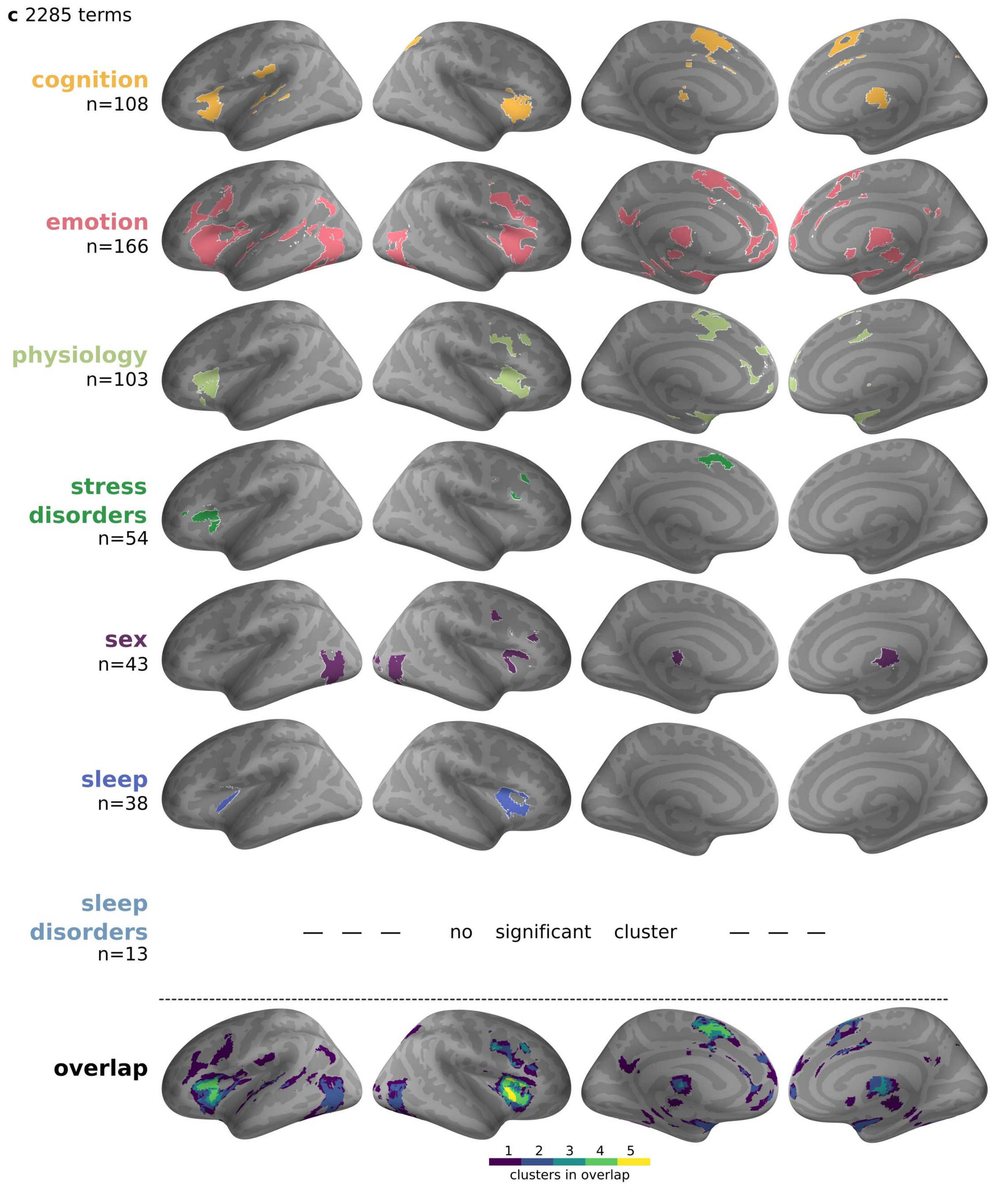


**Supplementary Figure 4. Brain-imaging results are stable for different numbers of terms in the semantic analysis.** Results of the neuroimaging meta-analysis across the number of terms (1500, 2000, 2285) included in the semantic analysis: **A.** 1500 terms, **B.** 2000 terms, **C.** 2285 terms. Across the number of terms used in the semantic analysis only small differences in the overlap are found. The cortical arousal network, composed of left and right anterior insula and preSMA is consistently found to be the overlap foci across all analyses.

**Supplementary Table 1.** Inclusion/Exclusion criteria **-** guidelines for term selection. We excluded terms related to, or terms that contained a word related to, the categories listed below.

| **Guidelines for exclusion (in quotes see literal examples)** | | |
|---|---|---|
| | literal example | exception/ comment |
| Non-specific general research related words: | | |
| Items describing statistical method, procedures and results like methods, descriptives, significance | “regression”, “SD”, “SVD”, “convolutional neural network”, “significant improvement” | due to ambiguity keep “neural network” |
| Items describing general study results | “behavioral results”,“Randomized trials” | |
| Items describing species & species subgroup | “rat model”, “human”, “C.elegans”,“heterosexual men”, “older adults”, “OSA patients” | “depressive patients” should be excluded but “depression” should be kept |
| Items describing subject pool and terms describing non-specific treatment of the experimental group | “control group”, “test group”, “treatment group” “treatment options”, “clinical variables”, “clinical features” | “Overweight group” should be excluded but “obesity” or “body mass” should be kept |
| Items describing change | “faster responses”, “treatment effect”,” increased result” “higher arousal” | KEEP terms describing the potential variable (“reaction time”, “FSFI scores”,” age effects”) or measurement tools (“fMRI”, “EEG” etc.)! |
| Items describing general experimental design | “twin study”, “crossover study”, “task demand” | KEEP terms describing experimental material, like task (ex. “stroop task”) or potential stimuli used (ex. “neutral scenes”, “video clips” etc.)! |

| | | |
|---|---|---|
| Items describing Editors, Journals, International/Scientific/Academic societies | “Sexual Medicine”, “Neurophysiology”, “International Society” | |
| **Others:** | | |
| Items that have little semantic significance or terms that don’t refer to any specific variables, for example | units (“micro gram”,” celsius degrees”),” last decade”, “large number”, “peak amplitude”, “frequency and duration”, “treatment approach”, “first night”, “greater risk”, “objective measures”, “Similar results”, “previous research”, “relative contribution”, “index score, “function scores” | importantly exclude “index score”, “function scores” as they’re not referring to any specific variable but keep “FSFI score” or “arousal index” as they refer to specific variables |
| Countries or other Geographical locations | Hong Kong, USA etc, | |

**Supplementary Table 2.** For each semantic community, the five terms with the largest number of connections within the community and their inside degree centrality.

| | Term, Degree inside |
|---|---|
| Cognition | Reaction time, 89<br>Choice reaction, 37<br>Cognitive performance, 36<br>Choice reaction time, 34<br>Task performance, 32 |
| Emotion | Emotional stimuli, 88<br>Neutral pictures, 54<br>Emotional pictures, 52<br>Unpleasant pictures, 44<br>Negative picture, 43 |
| Physiology | Heart rate, 203<br>Blood pressure, 131<br>Rate variability, 85<br>Hear rate variability, 85<br>Cardio-vascular response, 80 |
| Stress disorders | Stress disorder, 81<br>PTSD symptom, 68<br>Post-traumatic stress, 66<br>Post-traumatic stress disorder, 65<br>Trauma exposure, 50 |
| Sex | Sexual dysfunction, 95<br>Female sexual, 95<br>Sexual desire, 84<br>Sexual function, 84<br>Female sexual dysfunction, 78 |
| Sleep | Rapid eye, 117<br>Rapid eye movement, 116<br>Eye movement, 116<br>REM sleep, 96<br>nREM sleep, 96 |
| Sleep disorders | Obstructive sleep, 94<br>Sleep apnea, 91<br>Obstructive Sleep Apnea, 86<br>Apnea hypopnea, 73<br>Hypopnea index, 70 |

**Supplementary Table 3.** Dictionary of terms used to find physiological measures in the texts.

| **measure** | **Terms searched for** |
|---|---|
| heart | ['heart-rate', 'heart rate', 'hrv', 'interbeat interval', 'IBI', 'inter-beat interval', 'cardiac rate', 'cardiac response'] |
| skin conductance | ['skin conductance',’skin resistance’, 'electrodermal', 'galvanic skin response', 'SCR', 'GSR', 'EDA'] |
| pupil | ['pupil', 'pupillography', 'pupillometry', 'oculography', ‘oculometry’, ’eye tracking’, ‘eye-tracking’] |
| respiration | ['respiration', 'respiratory', 'breathing', 'inhalation', 'exhalation', 'ventilation', 'ventilatory'] |
| blood pressure | ['blood pressure', ’systolic pressure”, 'diastolic pressure'] |
